# Inhibition of SARS-CoV-2 infection in human cardiomyocytes by targeting the Sigma-1 receptor disrupts cytoskeleton architecture and contractility

**DOI:** 10.1101/2021.02.20.432092

**Authors:** José Alexandre Salerno, Thayana Torquato, Jairo R. Temerozo, Livia Goto-Silva, Mayara Mendes, Carolina Q. Sacramento, Natalia Fintelman-Rodrigues, Gabriela Vitoria, Leticia Souza, Isis Ornelas, Carla Veríssimo, Karina Karmirian, Carolina Pedrosa, Suelen da Silva Gomes Dias, Vinicius Cardoso Soares, Luiz Guilherme HS Aragão, Teresa Puig-Pijuan, Vinícius W. Salazar, Rafael Dariolli, Diogo Biagi, Daniel Rodrigues Furtado, Helena L. Borges, Patrícia Bozza, Marília Zaluar Guimarães, Thiago Moreno L. Souza, Stevens K. Rehen

**Author notes:** These authors contributed equally to this work (José Alexandre Salerno, Thayana Torquato). Corresponding author: Stevens Rehen.

## Abstract

Heart dysfunction, represented by conditions such as myocarditis and arrhythmia, has been reported in COVID-19 patients. Therapeutic strategies focused on the cardiovascular system, however, remain scarce. The Sigma-1 receptor (S1R) has been recently proposed as a therapeutic target because its inhibition reduces SARS-CoV-2 replication. To investigate the role of S1R in SARS-CoV-2 infection in the heart, we used human cardiomyocytes derived from induced pluripotent stem cells (hiPSC-CM) as an experimental model. Here we show that the S1R antagonist NE-100 decreases SARS-CoV-2 infection and viral replication in hiPSC-CMs. Also, NE-100 reduces cytokine release and cell death associated with infection. Because S1R is involved in cardiac physiology, we investigated the effects of NE-100 in cardiomyocyte morphology and function. We show that NE-100 compromises cytoskeleton integrity and reduces beating frequency, causing contractile impairment. These results show that targeting S1R to challenge SARS-CoV-2 infection may be a useful therapeutic strategy but its detrimental effects *in vivo* on cardiac function should not be ignored.

## 1. INTRODUCTION

COVID-19 is an airborne infectious disease caused by the Severe Acute Respiratory Syndrome Coronavirus 2 (SARS-CoV-2). Since the first patients were diagnosed with COVID-19, myocardial injury following SARS-CoV-2 infection has been reported (C. Huang et al., 2020). Serious lung infection leading to respiratory failure is the main cause of death (Gnecchi et al., 2020; L. Huang et al., 2020; Vincent and Taccone, 2020). However, acute myocardial injury is also ranked as a cause of SARS-CoV-2 infection-related fatalities; Shi and collaborators have shown that the in-hospital mortality was increased (of 51.2%) in comparison to cases without cardiac injury (of 4.5%) (Shi et al., 2020). Cardiovascular comorbidities are related to worse outcomes and, together with diabetes, are the most current chronic conditions among hospitalized COVID-19 patients (B. Li et al., 2020; Yang et al., 2020).

SARS-CoV-2 was reported in the myocardial tissue (Lindner et al., 2020; Yao et al., 2020). Whereas some autopsy studies suggested that SARS-CoV-2 infects cardiomyocytes (Bulfamante et al., 2020), others indicated that the most likely targets are interstitial or invading cells in the myocardium (Hoffmann et al., 2020; Lindner et al., 2020). Therefore, it is not a consensus whether the myocardial disease is a direct result of SARS-CoV-2 damage to cardiomyocytes or secondary to systemic events.

Angiotensin-converting enzyme 2 (ACE2) is reported as the main receptor mediating SARS-CoV-2 cell entry (Scialo et al., 2020). ACE2 is expressed in many organs including the heart, where it plays a pivotal role in cardiovascular function (Li, 2016; Lu et al., 2020; Patel et al., 2016; Shang et al., 2020). The number of ACE2-positive cells and the expression of ACE2 itself are increased in failing hearts, which could facilitate SARS-CoV-2 infection (Nicin et al., 2020; Xu et al., 2020). These observations suggest an interplay between the cardiovascular system and susceptibility to infection.

Analyzing the human embryonic heart, Yang and collaborators found that ACE2 is enriched in the cardiomyocyte subset of cells and argue that priming of the SARS-CoV-2 Spike protein (SP) probably occurs through host cell cathepsins B and L (CTSB and CTSL) (Yang et al., 2021). Following ACE2 receptor attachment, SARS-CoV-2 utilizes the CTSL-dependent endolysosomal route in human induced pluripotent stem cell-derived cardiomyocytes (hiPSC-CMs) (Pérez-Bermejo et al., 2020). The hiPSC-CMs reproduce many key features of human myocardial cells and have been recognized as useful tools to address SARS-CoV-2 heart infection *in vitro* and to test drugs that may eventually prevent cardiac susceptibility (Choi et al., 2020; Marchiano et al., 2020; Pérez-Bermejo et al., 2020; Sharma et al., 2020).

Several compounds with antiviral activity have been proposed against SARS-CoV-2 infection, especially from drug repurposing studies (Lovato et al., 2020; Reznikov et al., 2020; Vela, 2020). Negative modulators of the Sigma-1 receptor (S1R) gained considerable attention in SARS-CoV-2 infection because of its interaction with non-structural protein 6 (NSP6) from SARS-CoV-2 (Gordon et al., 2020). S1R binds ligands with very diverse structures from a wide spectrum of compounds and small molecules (Su et al., 2010; Tesei et al., 2018). Experimental and approved drugs that bind Sigma receptors with mild to high affinity, even as an off-target, have been considered to prevent or treat COVID-19 (Reznikov et al., 2020; Vela, 2020). Several of these compounds that inhibit S1R function have antiviral activity, including against other coronaviruses, but cardiotoxicity and induction of arrhythmias were also reported (Chen et al., 2006; Page et al., 2016).

On the other hand, apart from antiviral activity and participation in cardiac function, S1R activation has also drawn attention for its potential anti-inflammatory properties that could control the hyperinflammatory damage that follows severe SARS-CoV-2 infection (Troncone, 2020). Recently, a randomized placebo-controlled clinical trial showed that the S1R agonist fluvoxamine prevents clinical deterioration of symptomatic COVID-19 patients (Lenze et al., 2020). Indeed, S1R activity is frequently described as an inhibitory pathway of cytokine production via inositol-requiring enzyme 1-alpha (IRE1-α)(Rosen et al., 2019a). As a result of this downstream signaling, S1R stimulation was demonstrated to protect mice from septic shock. As current evidence shows that the outpouring of inflammatory cytokines at the late stages of critical COVID-19 cases has a clear correlation with severity of symptoms and poor prognosis (X. Li et al., 2020), is of utmost relevance to consider both antiviral and anti-inflammatory approaches for potential drug candidates.

The modulation of S1R is in the spotlight of alternative pharmacological approaches in COVID-19 but remains to be explored. In this study, we investigated the role of S1R in SARS-CoV-2 infection of hiPSC-CMs. Inhibition of S1R reduced SARS-CoV-2 infection and viral replication, preventing infection-associated cell death and cytokine release. On the other hand, it led to aberrant changes in the cytoskeleton and impaired cellular contraction. These results suggest that targeting S1R as a strategy against COVID-19 should be considered carefully regarding possible adverse cardiac outcomes.

## 2. RESULTS

### 2.1 Human induced pluripotent stem cell-derived cardiomyocytes (hiPSC-CMs) express the cardiac-specific troponin T and S1R

Human cardiomyocytes were differentiated from induced pluripotent stem cells (iPSCs) according to established protocols(Cruvinel et al., 2020). Differentiation to cardiomyocytes resulted in a cell population with low expression of the pluripotency marker OCT-4 (0.8% ± 0.4%) and most cells expressing the specific cardiac muscle marker troponin T (cTnT) (88.4% ± 8.4%), as assessed by flow cytometry (**Figure 1A**). The presence of cTnT was confirmed by immunostaining and cytoskeleton cell morphology can be visualized by F-actin staining (**Figure 1B**). Sigma receptors are known to be expressed in rat heart cells, especially the S1R subtype(Novakova et al., 1995). Here, we showed S1R expression in human cardiomyocytes by western blot (**Figure 1C**) and observed by immunostaining that S1R is widely spread within cells with an enhanced signal in the perinuclear region and the nuclei (**Figure 1D**). This is consistent with the cellular distribution of S1R in other cell types from rodents (Hayashi and Su, 2003).

**Figure 1.**
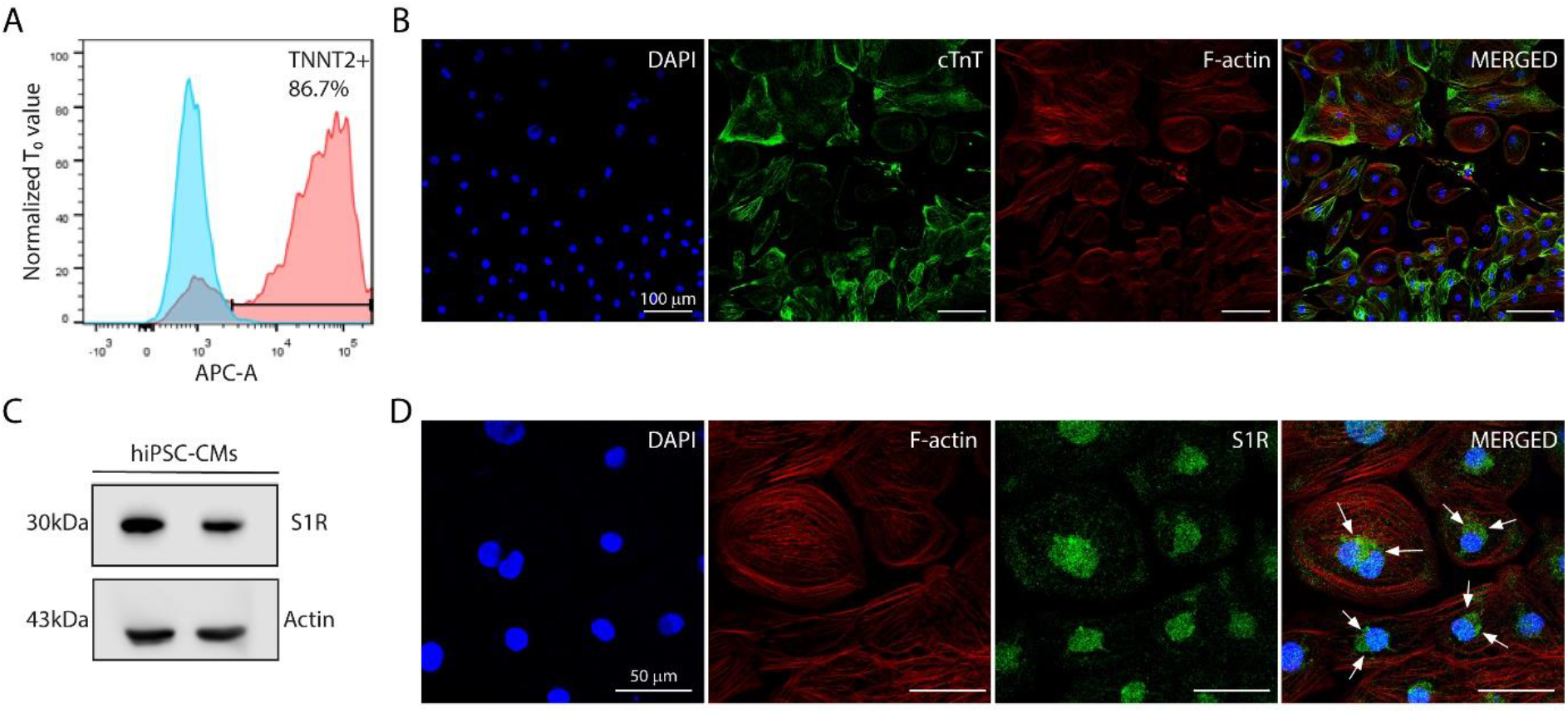
Human induced pluripotent stem cell-derived cardiomyocytes (hiPSC-CMs) express the specific cardiac Troponin T (cTnT) marker and the Sigma-1 receptor (S1R). **(A)** Flow cytometry data representing the expression of cardiac-specific marker troponin T (cTnT/*TNNT2*), confirming highly efficient differentiation into cardiomyocytes. **(B)** Immunocytochemistry for cTnT and phalloidin staining for filamentous actin (F-actin). Anti-cTnT (green); phalloidin (red) and nuclei (blue); scale bar = 100 μm. **(C)** Western blotting of S1R in protein extracts of control hiPSC-CMs performed in duplicate. Actin was used as a loading control. **(D)** Immunocytochemistry for S1R in hiPSC-CMs. Nuclear and perinuclear regions (white arrows). S1R (green); phalloidin (red) and nuclei (blue); scale bar = 50 μm.

### 2.2 Inhibition of S1R reduces SARS-CoV-2 infection and replication in human cardiomyocytes and prevents cell death

To inhibit S1R in hiPSC-CMs, cells were treated with the high-affinity S1R antagonist NE-100, which has a very low affinity for other receptors such as dopamine, serotonin, and phencyclidine receptors (Okuyama et al., 1993). NE-100 does not show toxicity at concentrations ranging from 10 nM to 10 μM after 72 h (**Supplemental Figure 1 A**). Also, hiPSC-CMs treated with 1 μM NE-100 for 48 h and 72 h showed no significant changes in the number of pyknotic nuclei, which represents an irreversible chromatin-condensed nuclear state characteristic of cell death (**Supplemental Figure 1 B**).

Following 24 h pretreatment with 1 μM NE-100, hiPSC-CMs were infected with SARS-CoV-2 and the infection rate was evaluated at 48 hours post-infection (h.p.i). Immunofluorescence using convalescent serum (CS) from a recovered COVID-19 patient showed that 57.3% (± 11.1%) of cells were infected at 48 h.p.i (**Figure 2A**). We confirmed that CS-staining mostly co-localizes with the specific SARS-CoV-2 Spike Protein (SP) (**Supplemental Figure 2**). Importantly, S1R inhibition reduced the percentage of infected hiPSC-CMs to 35.8% (± 2.5%) (**Figure 2A and B**).

**Figure 2.**
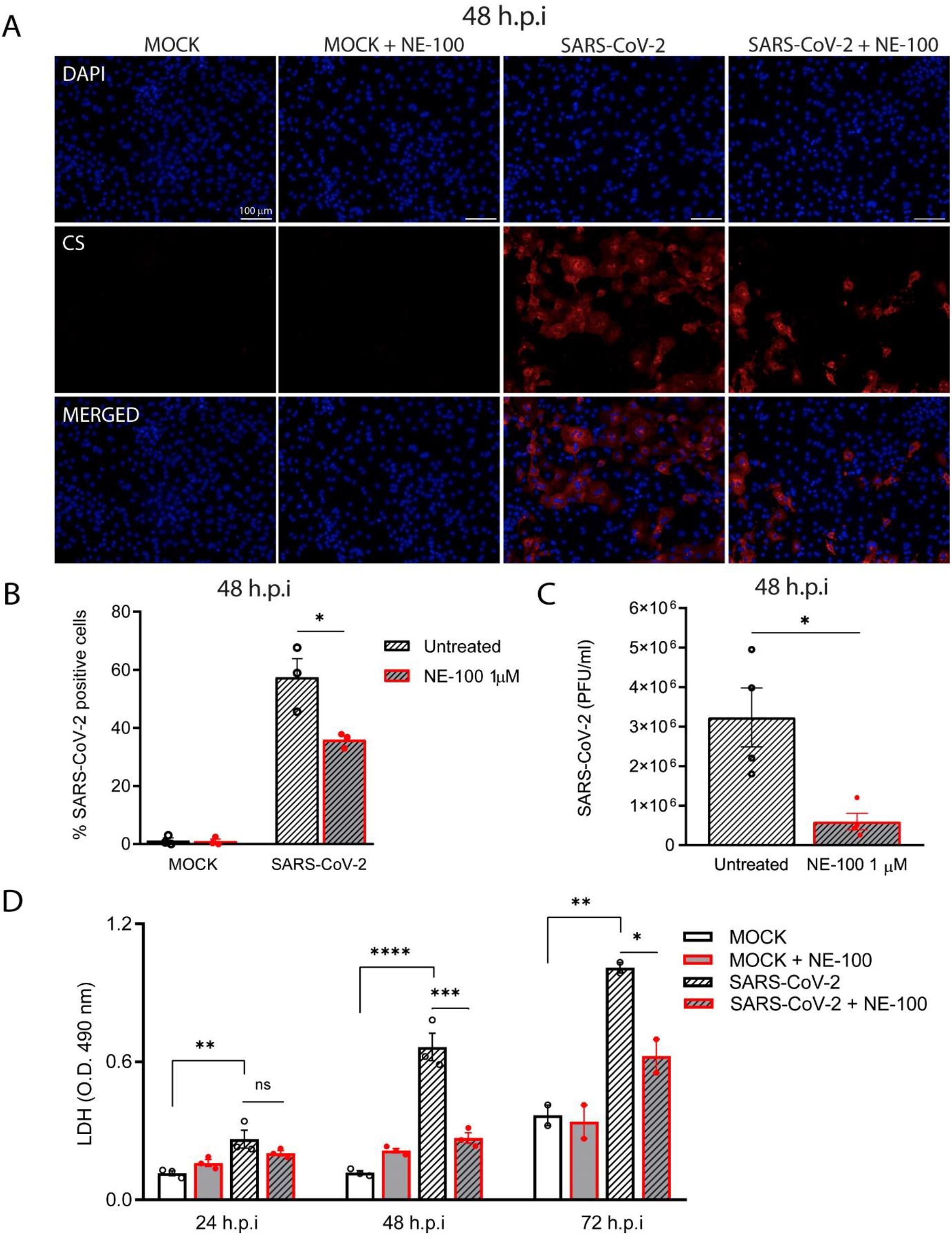
S1R inhibition reduces SARS-CoV-2 infection, replication and cytotoxicity in hiPSC-CMs. **(A)** hiPSC-CMs were pre-treated for 24 h with 1 μM NE-100 and infected with SARS-CoV-2 at multiplicity of infection (MOI) of 0.1. Cells were evaluated at 48 hours post-infection (h.p.i). Immunostainings show infected hiPSC-CMs positively stained for convalescent serum (CS) in red and no signal detection in mock conditions; scale bar = 100 μm. **(B)** The percentage of infected hiPSC-CMs was assessed by quantification of CS positive cells in SARS-CoV-2-infected and mock-infected conditions exposed or not to S1R high-affinity antagonist NE-100. **(C)** Plaque forming units assay for the supernatants of the SARS-CoV-2 infected hiPSC-CMs. **(D)** Cell death was measured in the supernatant by LDH activity at 24, 48 and 72 h.p.i. Data are represented as the mean ± S.E.M obtained in at least three independent experiments. *p < 0.05, **p < 0.01, ***p < 0.001, **** p < 0.0001.

Infection of hiPSC-CMs with SARS-CoV-2 led to the production of infectious virions as shown in **Figure 2C**. Exposure to NE-100 significantly diminished viral yield, with an average reduction of 82% at 48 h.p.i (**Figure 2C**). SARS-CoV-2 infection was shown to induce cytopathic features in hiPSC-CMs, mostly related to myofibrillar disruption and sarcomeric fragmentation, as described by Pérez-Bermejo and collaborators (Pérez-Bermejo et al., 2020). Likewise, we detected a pattern of fragmentation at 48 h.p.i, as shown in **Supplemental Figure 3** with cardiac TnT staining.

Lactate dehydrogenase (LDH) release was evaluated in hiPSC-CM cultures and confirmed that SARS-CoV-2 infection causes cardiomyocyte death, as previously reported (Sharma et al., 2020). At 24, 48 and 72 h.p.i, LDH levels in cell supernatants were elevated in SARS-CoV-2-infected conditions when compared to mock controls by 2.3, 5.6 and 2.7-fold respectively (**Figure 2D**). NE-100 significantly decreases LDH leakage at 48 and 72 h.p.i (**Figure 2D**). These results suggest that the inhibition of S1R may prevent hiPSC-CMs death by decreasing susceptibility to infection.

Since S1R is engaged in ER protein synthesis and acts as chaperone for proteins translocating to cell surface (Vela, 2020), we investigated whether S1R inhibition could be related to modifications of the host cell receptor for viral entry. To that end, the expression of ACE2 was evaluated after a 24 h treatment with 1 μM of S1R antagonist NE-100. We observed a reduction in the levels of ACE2 mRNA, not statistically significant (P=0.0790) (**Supplemental Figure 4A**). Western blot analysis of ACE2 after NE-100 treatment did not show differences at the protein level (**Supplemental Figures 4B and C**). These data suggest that the inhibition of SARS-CoV-2 infection in hiPSC-CMs can occur through mechanisms other than a reduction in the availability of ACE2 under the above described NE-100 exposure.

**Figure 4.**
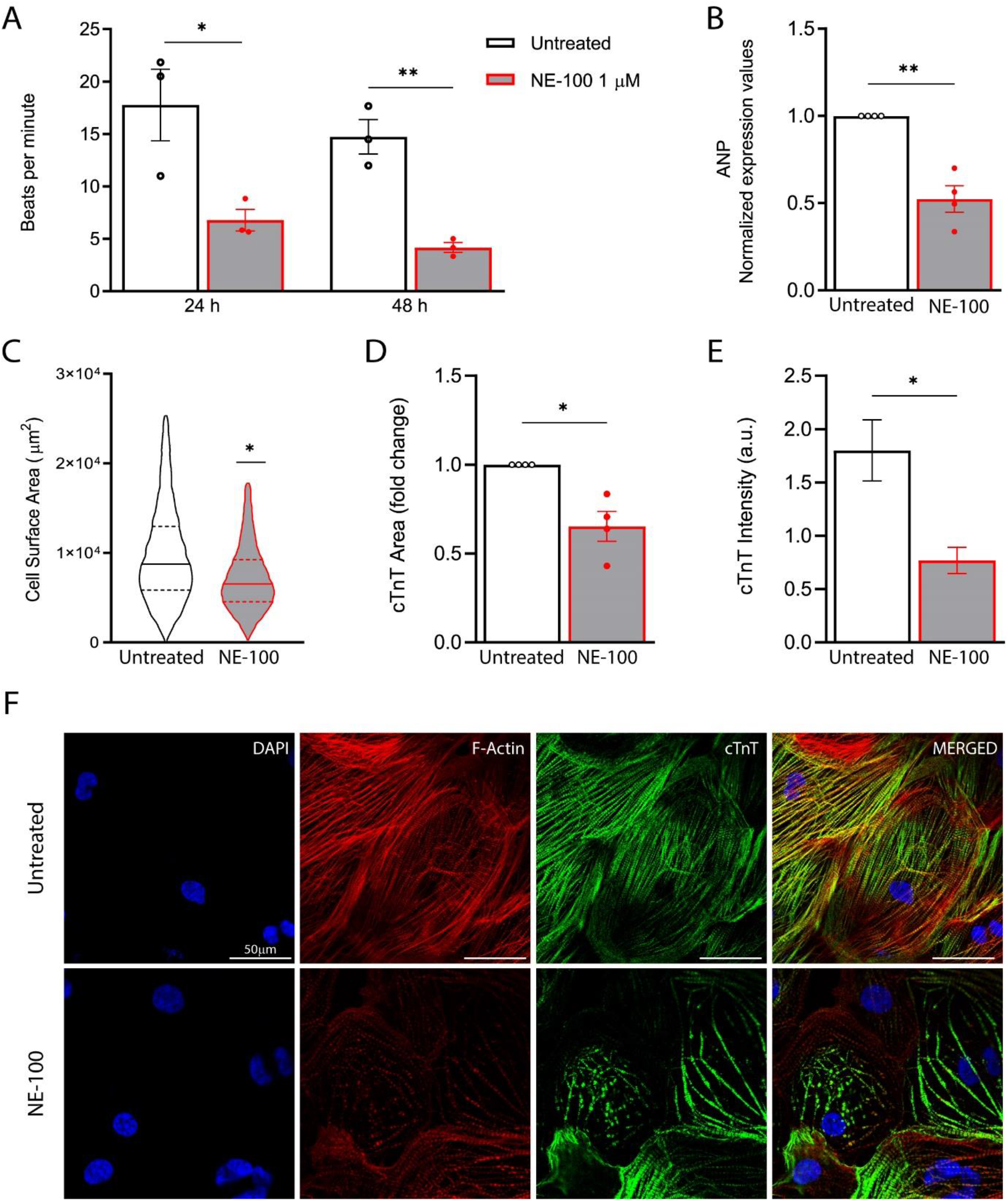
NE-100 decreases beating frequency and disrupts cytoskeleton integrity in hiPSC-CMs. **(A)** Average of beats per minute analyzed after exposure to NE-100, from three independent experiments. **(B)** Real-time PCR shows decreased levels of transcript content for ANP after S1R inhibition. **(C)** Cell area was quantified by F-actin staining and shows a decrease in cell body sizes after exposure to NE-100 for 48 h. **(D and E)** Quantification of cTnT immunoreactive area and intensity, normalized by the total number of cells per field; values are expressed relative to untreated controls. **(F)** Confocal images show in more detail the disruption of F-actin and cTnT organization. Scale bar = 50 μm. Data is presented as the average ± S.E.M from at least three independent assays. *p<0.05; **p<0.01

### 2.3 S1R inhibitor NE-100 attenuates cytokine release in SARS-CoV-2 infected hiPSC-CMs

Cell death is probably the final denouement to hiPSC-CMs at 72 h.p.i (Sharma et al., 2020). To investigate early events before cell death, we analyzed the levels of some of the main cytokines associated with COVID-19 at 24 and 48 h.p.i. We found that SARS-CoV-2 infection increased the release of interleukin (IL)-6 when compared to control. NE-100 decreased the release of interleukin (IL-6) at 24 h.p.i (**Figure 3A**). At 48 h.p.i, IL-6 increased 4-fold in comparison to control. Similarly, NE-100 decreased the release of IL-6 (**Figure 3B**).

**Figure 3.**
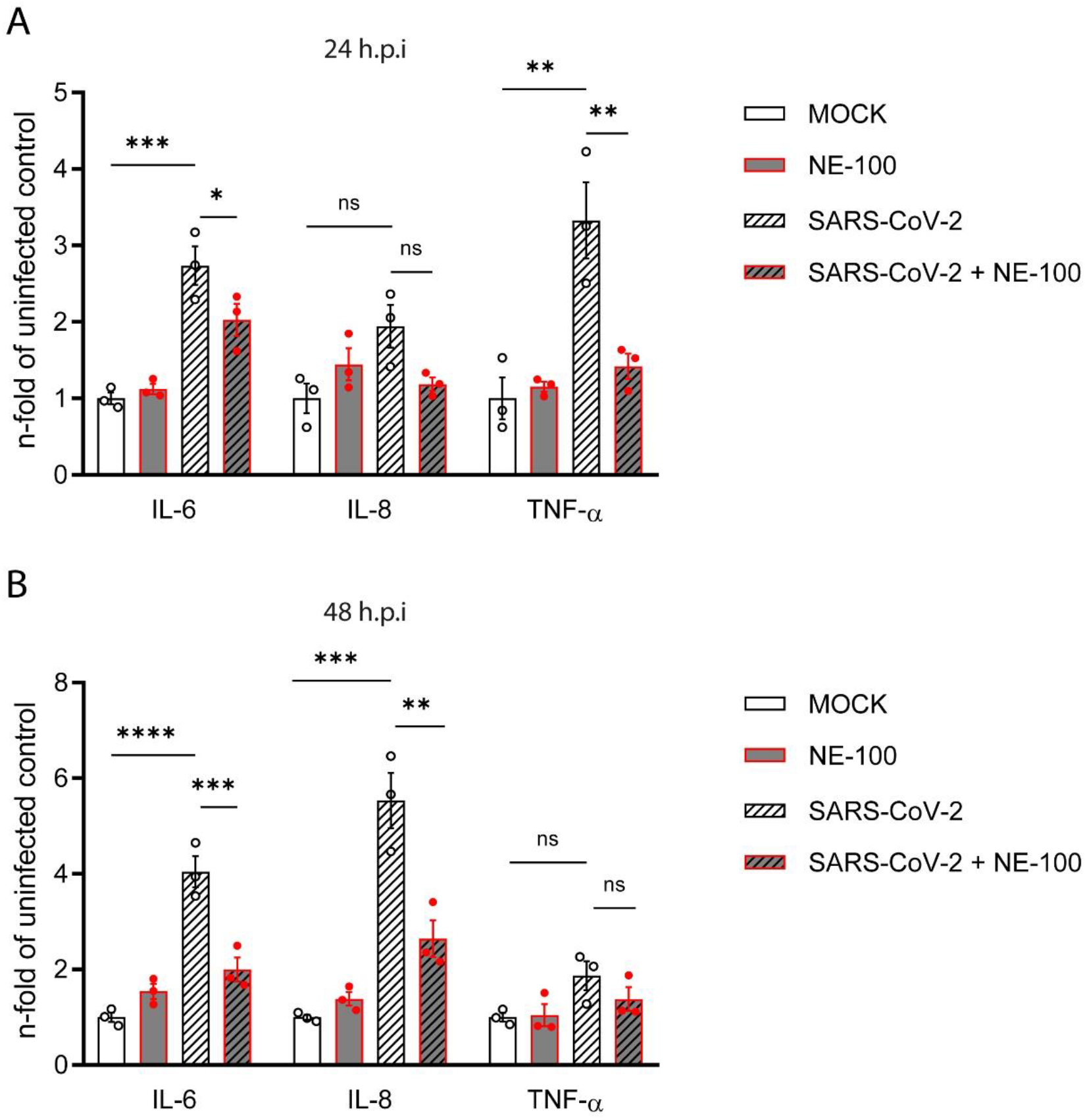
NE-100 decreases cytokine release that follows SARS-CoV-2 infection in hiPSC-CMs. **(A and B)** hiPSC-CMs were pre-treated or not with NE-100 and infected with SARS-CoV-2. Supernatants were analyzed by ELISA for IL-6, IL-8 and TNF-α at 24 h.p.i (A) and 48 h.p.i (B). Data represent mean ± SEM from three independent experiments. *p < 0.05, **p < 0.01, ***p < 0.001, **** p < 0.0001.

No distinctly clear differences in the IL-8 secretion pattern were apparent at 24 h.p.i (**Figure 3A**). However, at 48 h.p.i, SARS-CoV-2 increased IL-8 amount by 5.5-fold relative to mock condition (**Figure 3B**). Comparably to the modulation of IL-6 levels, NE-100 robustly decreased IL-8 secretion (**Figure 3B**). Tumor necrosis factor-alpha (TNF-α) was also measured and found increased only at 24 h.p.i (3.3-fold), when the NE-100 treatment restored secretion to similar levels of control condition (**Figure 3A**), while no significant changes were observed at 48 h.p.i (**Figure 3B**).

### 2.4 S1R inhibition disrupts cytoskeleton architecture and contractility of human cardiomyocytes

One of the main features displayed by cultured cardiomyocytes is their ability to perform spontaneous contractions (Belostotskaya and Golovanova, 2014). In that matter, hiPSC-CMs were proved to be a suitable model to evaluate drug-induced changes in contractility (Niehoff et al., 2019; Pointon et al., 2015). We observed a decrease from 17.7 (± 5.9) beats per minute (bpm) to 6.7 (± 1.8) bpm and from 14.7 (± 2.8) bpm to 4.1 (± 0.8) bpm after 24 h and 48 h of NE-100 exposure, respectively (**Figure 4A and Supplemental videos**). These results suggest that the inhibition of S1R leads to contractile impairment and a putative functional collapse.

S1R plays a cardioprotective role during maladaptive cardiac remodeling and the anti-hypertrophic properties of S1R agonists have been extensively described (Hirano et al., 2014; Tagashira et al., 2013; Tagashira and Fukunaga, 2012). Therefore, we also investigated if the S1R antagonist NE-100 alone could induce a hypertrophic response that could lead to contractile machinery overload and beating frequency reduction. To that end, we evaluated gene expression upon S1R inhibition, measuring mRNA levels of the atrial natriuretic peptide (ANP), which is one of the main transcripts of the fetal program for cardiac growth. ANP overexpression is frequently correlated to cardiac hypertrophy and NE-100 decreased mRNA expression of ANP by 1.49-fold after 24 h (**Figure 4B**). Next, we evaluated cell size and no increase in the average cell surface area of hiPSC-CMs was observed by F-actin staining. In fact, NE-100 exposure decreased the average cell surface area after 48 h (**Figure 4C**), which is a phenotype opposite to the expected for cardiomyocyte hypertrophic responses *in vitro* (Watkins et al., 2012).

We then hypothesized that the reduction in beating frequency could be related to cell shrinking and attributed to morphological changes induced by S1R inhibition. Analysis of cardiac troponin (cTnT) staining revealed a significant decrease both in cTnT immunoreactive area and fluorescence intensity of hiPSC-CMs exposed to NE-100 when compared to untreated controls (**Figure 4D and E, respectively**). These results are probably attributable also to a decrease in cellular cTnT content rather than exclusively to a change in cell area, indicating a noticeable loss of troponin. In addition, the integrity of cytoskeletal fibers was grossly affected, as shown in confocal fluorescence images (**Figure 4F**).

Since there are changes in the cytoarchitecture, but not in cell toxicity or hypertrophy induction, we suggest that cytoskeleton disruption is the most likely cause for contractile collapse of hiPSC-CMs induced by S1R inhibition.

## 3. DISCUSSION

In this study, we show that hiPSC-CMs are permissive to a SARS-CoV-2 productive infection, corroborating previous results and validating this model to study COVID-19 cardiac pathology *in vitro* (Bojkova et al., 2020; Bulfamante et al., 2020; Choi et al., 2020; Marchiano et al., 2020; Pérez-Bermejo et al., 2020; Sharma et al., 2020). Cell-based anti-SARS-CoV-2 drug screenings have been widely performed and the hiPSC-CMs approach is improving rapidly so that it can support the prediction of cardiac side effects at early preclinical steps of drug development (Chaudhari et al., 2016; Mirabelli et al., 2020; Yiangou et al., 2020).

We described that S1R inhibition with the high-affinity antagonist NE-100 decreases the number of SARS-CoV-2-infected hiPSC-CMs by 21.5% and viral yield by 82% at 48h.p.i. Sharma and colleagues previously described increased cell death through apoptosis in hiPSC-CMs after SARS-CoV-2 infection (Sharma et al., 2020). Here, we confirmed the cytotoxic effect of the virus providing evidence to support that direct damage to cardiomyocytes follows SARS-CoV-2 infection and could be related to cardiac injury in COVID-19.

Broader time point analyses are still needed to rule out an ACE2-dependent inhibition of infection by S1R antagonism. However, there is no description of direct interaction between S1R and host proteins involved in SARS-CoV-2 attachment or viral envelope proteins (Vela, 2020). Moreover, the assumption of a blockage mechanism after cell entry is in agreement with the SARS-CoV-2–human protein interaction map based on proteomics analysis, whereupon SARS-CoV-2 NSP6, located at the ER, interacts directly with human S1R (Gordon et al., 2020). During the replication of mammalian coronaviruses, this viral protein orchestrates vesicle trafficking and regulates ER remodeling (Cottam et al., 2011; J Alsaadi and Jones, 2019). This close relationship with S1R could be related to its activity in the rearrangement of endomembrane compartments and trafficking to favor steps of the replication of SARS-CoV-2 (Gordon et al., 2020; Santerre et al., 2020).

Gordon and colleagues assessed the antiviral activity of sigma receptor ligands in Vero E6 cells, which are highly permissive to infection. The authors described that those with antagonist action reduced viral infectivity by targeting replication (Gordon et al., 2020). Moreover, they reported that the S1R agonist dextromethorphan had proviral activity. Herein, we provide evidence to support that this is also true for human cardiac cells *in vitro*. Interestingly, compounds believed to protect from SARS-CoV-2 infection that are currently undergoing clinical trials, such as chloroquine (phase 2 NCT04349371), have S1R as an off-target (Hirata et al., 2011).

S1R is critical for early steps of Hepatitis C Virus (HCV) RNA replication so that its downregulation decreases susceptibility to infection in the hepatocyte-derived carcinoma cell lineage Huh-7 (Friesland et al., 2013). A hallmark of these positive-sense single-stranded RNA viruses, such as HCV and SARS-CoV-2, is that replication occurs inside modified membranes derived from the ER, where S1R is mostly expressed (Miller and Krijnse-Locker, 2008; Mori et al., 2013; Ritzenthaler and Elamawi, 2006). Approved drugs with antiviral activity against HCV replication have been demonstrated to be potential repurposed compounds to fight SARS-CoV-2 infection *in vitro* (Sacramento et al., 2020). Notably, remdesivir, which is an efficient antiviral drug against HCV, was approved for emergency treatment of COVID-19 patients requiring hospitalization (Eastman et al., 2020). Remdesivir anti-SARS-CoV-2 activity in human cardiomyocytes was described by Choi and colleagues, together with a safety profile evaluation that stipulated considerable arrhythmogenic and cardiotoxic risk *in vitro*, raising the concern of drug-induced cardiotoxicity of repositioned pre-approved compounds to manage COVID-19 (Choi et al., 2020).

Aside from viral infection and replication, the immune-mediated mechanisms are a hallmark of COVID-19 pathology and associated cardiac deterioration that can be successfully modeled using hiPSC-CMs (Azkur et al., 2020; Dariolli et al., 2021). Along with monocytes and fibroblasts, cardiomyocytes are also an important source of cytokines during events such as heart failure (Aoyagi and Matsui, 2012). Here, we showed that IL-6, IL-8 and TNF-α release is stimulated in hiPSC-CMs in response to SARS-CoV-2 infection, complementing previous data on increased mRNA levels of these cytokines (Wong et al., 2020). Elevated levels of IL-6 are strongly correlated with cardiac damage and heart failure in rodent models (Janssen et al., 2005; Jug et al., 2009). Moreover, previous reports demonstrated that IL-6 produced in cardiomyocytes promotes inflammation in the heart by recruiting neutrophils (Youker et al., 1992). Interestingly, IL-8, which was also found increased, is a neutrophil chemotactic factor correlated with chronic heart failure, coronary heart disease and myocardial apoptosis (Akasaka et al., 2006; Nymo et al., 2014; Rothenbacher et al., 2006). Indeed, myocardial infiltration of neutrophils has been reported in the hearts of COVID-19 patients and is believed to be a key mechanism behind myocardial damage (Yao et al., 2020).

The consequences of infection to cardiomyocytes are just beginning to be unraveled *in vitro* and should help to better understand COVID-19 heart pathology and its crosstalk with systemic events of disease progression. Our results suggest that immunopathological mechanisms probably underlie myocardial injury following SARS-CoV-2 infection and that cardiomyocytes could also play a role in the excessive systemic release of cytokines observed in the late stages of COVID-19. Interestingly, S1R inhibition with NE-100 attenuated the cytokine release following SARS-CoV-2 infection. However, the anti-inflammatory role promoted by S1R activation in the control of septic shock and its immunomodulatory activity during infections by other respiratory viruses, such as H1N1 influenza virus, have been well described (Rosen et al., 2019b; Szabo et al., 2014; Zhou et al., 2019). Therefore, the effect observed in this work is more likely to be due to the inhibition of infection than a direct effect on cytokine production.

Although S1R antagonism was an efficient mechanism to diminish SARS-CoV-2 infection and associated cell-death and cytokine release, there are reasons to believe that such inhibition could be detrimental to cardiac physiology (Shinoda et al., 2016b, 2016a). Also, as COVID-19 patients are expected to present cardiac dysfunctions (Magadum and Kishore, 2020; Wu et al., 2020), the use of drugs with potential cardiac negative effects could be unfeasible as a therapeutic approach.

Notwithstanding, the cell survival rate was not affected, and neither the characteristic hypertrophic gene expression profile nor enlargement of cell body sizes was detected after NE-100 exposure. Our findings show that inhibition of S1R significantly reduced beating frequency in hiPSC-CMs and completely compromised cell morphology with a conspicuous perturbation of cytoskeletal architecture and decrease in cardiac troponin staining. Actin subunits are assembled into F-actin, which serves as a platform for troponin-tropomyosin binding for proper sarcomere function during contraction (Gunning et al., 2015). The integrity of actin and troponin fibers dictates the mechanical stability necessary for proper force distribution at sarcomere margins during contraction and appropriate anchorage between cytoskeletal proteins and extracellular matrix (Gunning et al., 2015). The disruption of cytoskeletal and sarcomeric proteins by a decrease in expression or anomalous arrangement could underlie the pathogenesis of cardiomyopathies and heart failure (Sequeira et al., 2014).

It could be argued that hiPSC-CMs are too immature to reliably predict heart response to NE-100. However, the S1R antagonist haloperidol was demonstrated to prolong QT interval, deplete ATP production and cause shortening of action potentials using cardiomyocytes from different sources, including adult rodents (Shinoda et al., 2016a; Stracina et al., 2015; Tarabová et al., 2009). These observations were consistent with the deterioration of heart function in animal models. Furthermore, S1R knockout mice exhibit maladaptive morphological cardiac remodeling and thereafter contractile dysfunction (Abdullah et al., 2018).

Further investigations are needed to determine whether NE-100 might induce cardiotoxicity *in vivo*. However, it is hypothesized that the direct interaction between S1R and the *human Ether‐à‐go‐go Related Gene* (hERG) voltage-gated potassium channel could underlie the blockage of hERG function by S1R ligands, causing delays in cardiac repolarization, impairment of rate adaptation and increased risk for drug-induced arrhythmia (Balasuriya et al., 2014; Corbera et al., 2006; Crottès et al., 2011; Eng et al., 2016; Hancox and Mitcheson, 2006; Morales-Lázaro et al., 2019; Witchel, 2011; Yu et al., 2015).

Besides cytoskeleton-sarcomere framework integrity, the contractile capacity of cardiomyocytes is related to calcium availability (Van der Velden et al., 2003). As S1R is a key regulator of intracellular calcium homeostasis, the possibility that blocking its function could be hampering proper calcium cytoplasmic availability and impairing contraction cannot be ruled out, since this was not evaluated at the present work. However, the involvement of calcium intracellular concentration in S1R function in cardiac cells is controversial. Despite the previously described interaction of S1R inhibitors with L-type calcium channels in rat cardiomyocytes, the sensitivity of the myofilaments to calcium ions was shown not to be changed, while beating frequency variations followed by asymmetrical contractions were observed after exposure to S1R antagonists (Ela et al., 1994; Tarabová et al., 2009).

S1R has been extensively studied regarding its vital role for physiological cardiac function, amelioration of ER stress, and protection against maladaptive hypertrophy that leads to heart failure (Bhuiyan and Fukunaga, 2009; Tagashira et al., 2013; Tagashira and Fukunaga, 2012) but the role of S1R in human cardiomyocytes remains poorly described. Here, we provide data about the involvement of S1R in SARS-CoV-2 replication. Our *in vitro* results suggest that the inhibition of S1R as a therapeutic strategy against COVID-19 should be further investigated and undergo well-balanced decision-making when translated to clinical application, due to the concern of possible drug-induced cardiac malfunction.

## 4. MATERIAL AND METHODS

### 4.1 hiPSC-CMs differentiation, purification, and maintenance

Fresh human iPSCs-derived cardiomyocytes were purchased from Pluricell (São Paulo, Brazil) and the protocol for cardiomyocyte differentiation is described in detail by Cruvinel and colleagues (Cruvinel et al., 2020). Briefly, hiPSCs were passed and replated to be maintained in E8 medium with 5 μM of Y-27632 (Cayman Chemical, USA) with daily changes of media. After 100% confluence was reached, cells were kept in RPMI supplemented with B27 without insulin (both from Thermo Fisher, USA) and with 4 μM CHIR99021 (Merck Millipore Sigma, USA) for one day. After 24 h, the medium was supplemented with 10 ng/ml BMP4 (R&D Systems, USA) for an additional 24 h. On day 2, fresh medium was supplemented with 2.5 μM KY2111 and XAV939 (both from Cayman Chemical, USA). From day 4, cells were cultivated with CDM3 media (RPMI supplemented with 213 μg/ml ascorbic acid (Sigma Aldrich, USA), 500 μg/ml Bovine Serum Albumin (BSA) and 2 μg/ml Plasmocin (InvivoGen, USA). On day 18, cells were passaged with TrypLE 10X and plated on a density of 0.9×10^4^/well or 1.8×10^4^/well onto 96 or 24-well plates and maintained in CDM3 media. Upon arrival, cardiomyocytes were allowed to regain contractility and maintained at 37°C in a humidified atmosphere with 5% CO2. Cardiomyocytes were used between days 30 to 40 of differentiation.

### 4.2 Chemicals

4-Methoxy-3-(2-phenylethoxy)-N,N-dipropylbenzeneethanamine hydrochloride (NE-100 hydrochloride) was purchased from Tocris (3133). Stock and work solutions were prepared using 100% dimethyl sulfoxide sterile-filtered (DMSO; D2650 - Sigma-Aldrich).

### 4.3 Flow Cytometry

Cardiomyocytes were plated on 6-well plates coated with GELTREX and cultivated for 7 days. After cell dissociation, cells were fixed with 1% paraformaldehyde (PFA), permeabilized with Triton 0.1% (Sigma Aldrich) and Saponin 0.1% (Sigma Aldrich), and stained with the antibodies anti-TNNT2 (1:2500; Thermo Fisher, MA5-12960) and anti-OCT4 (1:200, Thermo Fisher, MA5-14845). FC data was acquired using a Canto BD flow cytometer for each batch of differentiation and analyzed using the FlowJo Software considering 1%–2% of false-positive events.

### 4.4 SARS-CoV-2 propagation

SARS-CoV-2 was expanded in Vero E6 cells from an isolate obtained from a nasopharyngeal swab of a confirmed case in Rio de Janeiro, Brazil (GenBank accession no. MT710714). Viral isolation was performed after a single passage in cell culture in 150 cm^2^ flasks with high glucose DMEM plus 2% FBS. Observations for cytopathic effects were performed daily and peaked 4 to 5 days after infection. All procedures related to virus culture were handled in biosafety level 3 (BSL3) multi-user facilities according to WHO guidelines. Virus titers were determined as plaque-forming units (PFU/ml), and virus stocks were kept in −80°C ultra-low temperature freezers.

### 4.5 Infections and virus titration

Cardiomyocytes were infected with SARS-CoV-2 at MOI of 0.1 in CDM3 media without serum. After 1 hour, cells were washed and incubated with complete medium with treatments or not. For virus titration, monolayers of Vero E6 cells (2 × 10^4^ cell/well) in 96-well plates were infected with serial dilutions of supernatants containing SARS-CoV-2 for 1 hour at 37°C. Semi-solid high glucose DMEM medium containing 2% FSB and 2.4% carboxymethylcellulose was added and cultures were incubated for 3 days at 37°C. Then, the cells were fixed with 10% formalin for 2 h at room temperature. The cell monolayer was stained with 0.04% solution of crystal violet in 20% ethanol for 1 h. Plaque numbers were scored in at least 3 replicates per dilution by independent readers blinded to the experimental group and the virus titers were determined by plaque-forming units (PFU) per milliliter.

### 4.6 Immunocytochemistry and fluorescence image analysis

hiPSC-CMs grown on 96-well plates were fixed using 4% PFA solution (Sigma-Aldrich) for 1 h and stored at 4°C until further processing. Cells were washed with 1X PBS and then incubated with permeabilization/blocking solution (0.3% Triton X-100/3% bovine serum albumin) for 1 h. Primary antibodies were diluted in blocking solution and incubated at 4°C overnight, namely anti-SARS-CoV-2 convalescent serum from a recovered COVID-19 patient (1:1000); anti-SARS-CoV-2 spike protein monoclonal antibody (SP) (1:500, G632604 - Genetex); anti-cardiac troponin T (cTnT) (1:500, MA5-12960 - Invitrogen) and anti-Sigma1R B-5 (1:100, SC-137075 - Santa Cruz Biotechnology). The use of the convalescent serum from COVID-19 patients was approved by CAAE number: 30650420.4.1001.0008. Next, hiPSC-CMs were washed 3 times with PBS 1X and incubated with the secondary antibodies diluted in blocking solution: goat anti-Human Alexa Fluor 647 (1:400; A-21445 - Invitrogen) and goat anti-Mouse Alexa Fluor 488 (1:400; A-11001 - Invitrogen) for 1 h at room temperature. Actin filaments were stained with Alexa Fluor 568 phalloidin (1:10; A-12380 - Life Technologies) for 1 h. Nuclei were stained with 300 nM 4ʹ-6-diamino-2-phenylindole (DAPI) for 5 minutes and each well was mounted with two drops of 50% PBS-Glycerol before image acquisition.

For quantitative analysis, images were acquired using Operetta® High-Content Screening System (Perkin Elmer) with a 20x long working distance (WD) objective lens from at least 10 fields per well. For cell surface area measurement, images of F-actin stained cardiomyocytes were segmented using Cellpose and the area was measured using NIH ImageJ software (Stringer et al., 2021). For the other analyses, data were evaluated using the Columbus™ Image Data Storage and Analysis System (Perkin Elmer) for image segmentation and object detection. The fluorescence threshold was set to determine positive and negative cells for each marker. Representative immunostaining images were acquired on a Leica TCS-SP8 confocal microscope using a 63x oil-immersion objective lens.

### 4.7 Measurements of cell death and inflammatory mediators

Monolayers of hiPSC-CMs in 96-well plates (70-90% confluence) were allowed to regain contractility and then were treated with various concentrations of NE-100. Neutral red (N4638 - Sigma-Aldrich) solution was prepared using ultrapure water and centrifuged (600 g for 20 minutes) to remove precipitates of dye crystals. At 72 hours post-treatment, media was removed, cells were washed once with PBS 1x and 200 μL of neutral red working solution diluted in hiPSC-CMs medium was added to each well at a final concentration of 25 μg/ml. Cells were incubated for 3 h at 37°C to allow uptake of dye into viable cells. Thereafter, media was removed, and cells were washed with PBS 1x followed by the addition of 1% acetic acid-49% ethanol revealing solution. After additional 10 minutes at 37°C and gently shaking, the absorbance was measured in the microplate reader spectrophotometer Infinite® M200 PRO (Tecan) at a wavelength of 540 nm. Surviving hiPSC-CMs were estimated by the percentage relative to untreated condition (vehicle) using the mean of 6 technical replicates per experiment.

The levels of IL-6, IL-8 and TNF-α were quantified in the supernatants from uninfected and SARS-CoV-2-infected hiPSC-CMs by ELISA (R&D Systems), following manufacturer’s instructions. Control groups (MOCK and cells infected with Sars-Cov-2 only) were also analyzed in Aragao et al., 2021 (in preparation). The results were obtained as picograms per milliliter (pg/ml) and are expressed as fold-change relative to untreated uninfected control. Cell death of infected cardiomyocytes was determined according to the activity of lactate dehydrogenase (LDH) in the culture supernatants using a CytoTox® Kit (Promega, USA) according to the manufacturer’s instructions. Supernatants were centrifuged at 5,000 rpm for 1 minute, to remove cellular debris.

### 4.8 Gene expression analysis

Total RNA was isolated using TRIzol reagent from hiPSC-CMs pellets, according to the manufacturer’s recommendations (Thermo Fisher Scientific). After, total RNA was digested with DNase using DNase I, Amplification Grade, following the manufacturer’s instructions (Invitrogen, Thermo Fisher Scientific). DNase-treated RNA samples (1 μg) were converted to complementary DNA (cDNA) using the M-MLV enzyme (Thermo Fisher Scientific).

The reactions to determine gene expression were conducted in three replicates with a final reaction volume of 10 μL in MicroAmp Fast Optical 96 Well Reaction Plates (Thermo Fisher Scientific) containing 1X GoTaq qPCR Master Mix (Promega Corporation), 300 nM CXR Reference Dye, final concentration 200nM of each SYBR green designed primers for the following targets: Angiotensin I Converting Enzyme 2 (ACE2; forward: 5’-CGAAGCCGAAGACCTGTTCTA-3’; reverse: 5’-GGGCAAGTGTGGACTGTTCC-3’); Natriuretic Peptide A (ANP; forward: 5’-CAACGCAGACCTGATGGATTT-3’; reverse: 5’-AGCCCCCGCTTCTTCATTC-3’); and 10 ng of cDNA per reaction. Briefly, the reactions were performed on a StepOnePlus ™ Real-Time PCR System thermocycler (Applied Biosystems). Thermal cycling program comprised hold stage at 95°C for 3 min, followed by 40 cycling stages at 95°C for 15 sec, 57°C for 15 sec, 72°C for 15 sec and melt curve stage 95 °C, 15 sec; 60 °C, 1 min; 95 °C, 15 sec. The relative expression of the genes of interest (GOI) was normalized by endogenous control genes: Glyceraldehyde-3-phosphate Dehydrogenase (GAPDH; forward: 5’-GCCCTCAACGACCACTTTG-3’; reverse: 5’-CCACCACCCTGTTGCTGTAG-3’) and Hypoxanthine Phosphoribosyl transferase 1 (HPRT-1; forward 5’-CGTCGTGATTAGTGATGATGAACC-3’; reverse: 5’-AGAGGGCTACAATGTGATGGC-3’). qPCR data analysis was performed with the N_0_ method implemented in LinRegPCR v. 2020.0 software, which considers qPCR mean efficiencies estimated by the window-of-linearity method as proposed by Ramakers et al. (2003) and Ruijter et al. (2009) (Ramakers et al., 2003; Ruijter et al., 2009). N_0_ values were calculated in LinRegPCR using default parameters and the arithmetic mean of N_0_ values from GOI were normalized by taking its ratio to the N_0_ of the geometric mean of the endogenous control genes (REF) GAPDH and HRRT-1 (N_0_GOI/N_0_REF).

### 4.9 Protein expression

Media was completely removed from hiPSC-CMs plated on 24-well plates, and 40μL of sample buffer without bromophenol blue (62.5 mM Tris-HCl, pH 6.8, containing 10% glycerol, 2% SDS, and 5% 2-mercaptoethanol) was added to each well. The lysate was frozen at −80°C until further processing. Next, cell extracts were boiled at 95°C for 5 min, centrifuged 16,000 × g for 15 min at 4°C, and the supernatant was collected. Protein content was estimated using the Bio-Rad Protein Assay (#5000006, Bio-Rad). After the addition of bromophenol blue (0.02%), extract samples were separated by electrophoresis on an 8% SDS polyacrylamide gel and transferred to polyvinylidene difluoride (PVDF) membranes.

Western blotting was performed with minor modifications from the originally described (Towbin et al., 1989). Briefly, membranes were blocked in 5% non-fat milk diluted in Tris-Buffered Saline with 0.1% Tween-20 (TBS-T) for 1 h at room temperature. Membranes were then incubated overnight at 4°C with primary antibodies (anti-ACE2 (1:1000; MA5-32307 - Thermo Fisher), anti-Sigma-1 R (1:500; SC-137075 - Santa Cruz Biotechnology), anti-actin (1:2000, MAB1501, Millipore); or anti-GAPDH (1:5000; AM4300 -Thermo Fisher) diluted in TBS-T with 5% non-fat milk. Then, membranes were washed and incubated with peroxidase-conjugated antibodies (goat anti-Mouse IgG (H+L), HRP-conjugate (1:10,000, G21040 -Molecular Probes) and Goat anti-Rabbit IgG (H+L) HRP- conjugate (1:10,000, G21234 - Molecular Probes). The signals were developed using ECL Prime Western Blotting System (#GERPN2232, Sigma) for 5 minutes and chemiluminescence was detected with an Odyssey-FC System® (LI-COR Biosciences).

Stripping protocol was performed to break bonds between the antibodies and the transferred proteins in order to reuse membranes. Briefly, membranes were incubated for three cycles of 10 minutes in a stripping buffer (pH 2.2, 200 mM glycine, SDS 0.1% and 1% Tween-20). Then, the buffer was discarded, the membranes were washed for 5 minutes with PBS (three times) and 5 minutes with 0.1% TBS-T (three times). Next, membranes were blocked again and proceeded with the above-described steps.

Densitometry analysis was performed using ImageJ software Gel Analysis program and the values obtained represent the ratio of density between the immunodetected protein and the loading control (actin or GAPDH).

### 4.10 Beating frequency evaluation

hiPSC-CMs were plated on 96-well plates (1.8×10^4^ cells per well) and treated with NE-100 1μM or vehicle (DMSO) before incubating at 37°C. At 24- and 48-hours post-treatment, beating frequency was measured by manually counting the synchronous contractions of the monolayer for 60 seconds (beats per minute). Cells were observed using an EVOS cell imaging system (Thermo Fisher Scientific), in brightfield mode. Three wells at a time were counted and then the plates were replaced for 5-10 minutes at the incubator and allowed to regain contractility before resuming counting. Representative videos were recorded using Operetta® High-Content Screening System (Perkin Elmer) for each condition at baseline and 48 h after treatment with vehicle or NE-100

### 4.11 Statistical analysis

Data are presented as mean values, and error bars indicate the standard error of the mean (S.E.M) from at least three independent experiments. Immunostaining quantification values were analyzed using nested t-test. For the other data, comparisons between two groups were performed with unpaired two-tailed Student’s t-test with Welch’s correction and between three or more groups, one-way ANOVA with Holm-Sidak post-hoc. Prism v8.0 (GraphPad) was used for data analysis and graphics, where statistical significance was accepted at P<0.05. The P values are specified in figure legends and the symbols represent *p < 0.05, **p < 0.01, ***p < 0.001, ****p < 0.0001.

## Supporting information

Supplemental Video - Untreated baseline

Supplemental Video - Untreated 48 h

Supplemental Video - NE-100 baseline

