## Supplemental Figure 1 for "Inhibition of SARS-CoV-2 infection in human cardiomyocytes by targeting the Sigma-1 receptor disrupts cytoskeleton architecture and contractility"

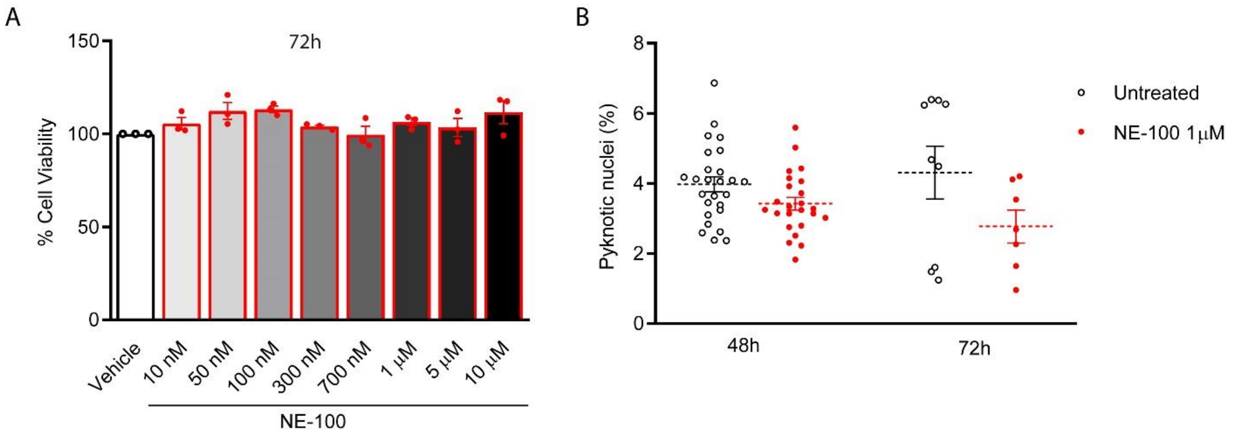


**Supplemental Figure 1. (A)** NE-100 does not induce cytotoxic effects in hiPSC-CMs. Neutral red cell viability assay for escalating NE-100 concentrations shows non-significant changes after 72 hours post-treatment. **(B)** Nuclear size analysis by DAPI staining shows the percentage of pyknotic nuclei in different conditions; Dots represent the percentage of each well analyzed in at least 2 independent experiments; non-significant.
