## Supplemental Figure 2 for "Inhibition of SARS-CoV-2 infection in human cardiomyocytes by targeting the Sigma-1 receptor disrupts cytoskeleton architecture and contractility"

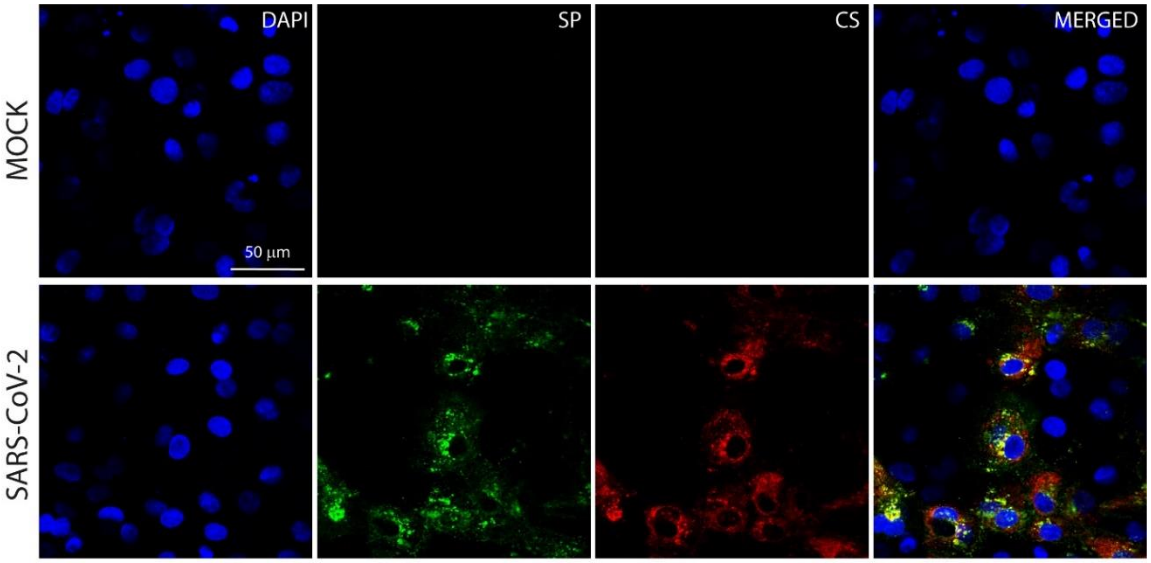


**Supplemental Figure 2.** Determination of the staining specificity of the convalescent serum from an in-house patient in comparison with the commercial antibody against the SARS-CoV-2 spike protein (SP). The CS showed robust and specific immunoreactive signals overlapped with the SP staining in SARS-CoV-2 infected cardiomyocyte cultures (MOI 0.1 at 48 h.p.i), while no such signal was detected in mock-infected condition. Scale bar = 50 µm.
