## Supplemental Figure 3 for "Inhibition of SARS-CoV-2 infection in human cardiomyocytes by targeting the Sigma-1 receptor disrupts cytoskeleton architecture and contractility"

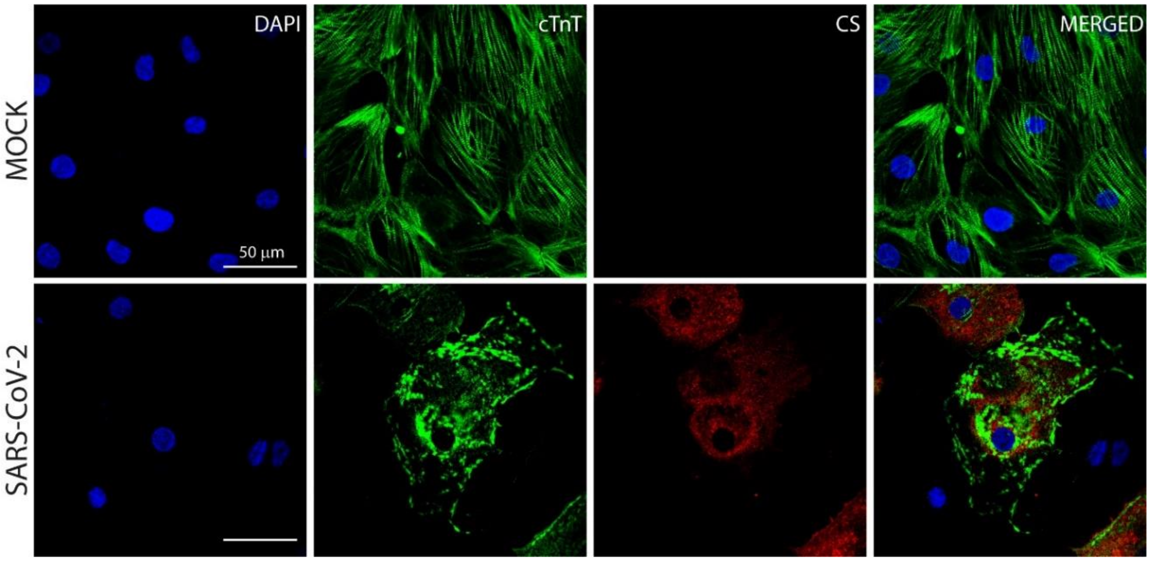


**Supplementary Figure 3.** Representative immunostaining images of cytoskeleton fragmentation of cardiomyocytes exposed to SARS-CoV-2 (MOI 0.1) for 48 h from 4 independent experiments. Anti-convalescent serum (CS) positive staining indicates the presence of SARS-CoV-2 in cells exhibiting Cardiac TnT discontinuity and disruption. cTnT (green), CS (red) and nuclei (blue). Scale bar = 50 µm.
