## Supplemental Figure 4 for "Inhibition of SARS-CoV-2 infection in human cardiomyocytes by targeting the Sigma-1 receptor disrupts cytoskeleton architecture and contractility"

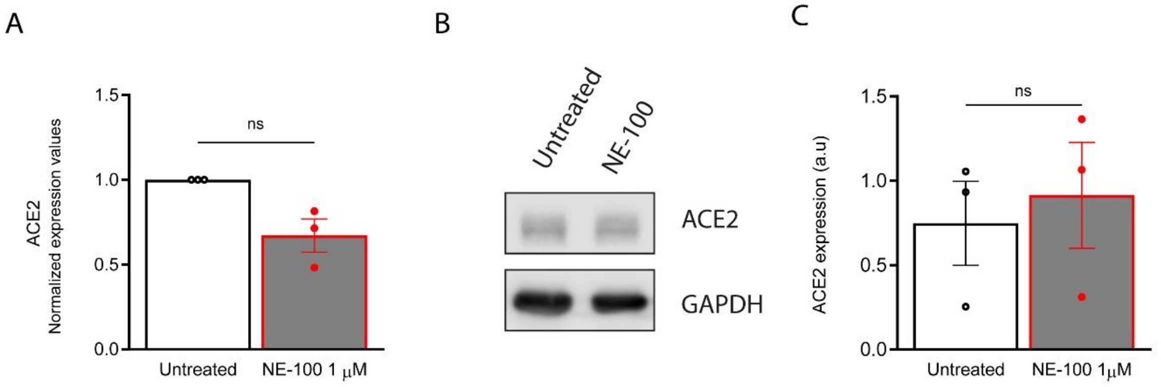


**Supplemental Figure 4. S1R inhibition does not alter ACE2 expression in hiPSC-CMs. (A)** Real-time PCR shows ACE2 mRNA levels. Expression values are normalized by endogenous control genes GAPDH and HPRT1 and are expressed as fold change relative to control (untreated) condition. **(B)** Western blots for ACE2 in protein extracts from untreated hiPSC-CMs or stimulated with 1µM of NE-100 for 24h. **(C)** Blots were quantified by densitometry and normalized by GAPDH expression. Data are presented as mean ± S.E.M of values from three independent experiments.
